# EWSR1 prevents the induction of aneuploidy by regulating the localization of Aurora B at inner centromere

**DOI:** 10.1101/2022.06.17.496636

**Authors:** Haeyoung Kim, Hyewon Park, Evan T. Schulz, Yoshiaki Azuma, Mizuki Azuma

## Abstract

*EWSR1 (Ewing sarcoma breakpoint region 1)* was originally identified as a part of an aberrant *EWSR1/FLI1* fusion gene in Ewing sarcoma, the second most common pediatric bone cancer. Due to formation of the *EWSR1/FLI1* fusion gene in the tumor genome, the cell loses one wild type *EWSR1* allele. Our previous study demonstrated that the loss of *ewsr1a* (homologue of human *EWSR1)* in zebrafish leads to the high incidence of mitotic dysfunction, of aneuploidy, and of tumorigenesis in the *tp53* mutant background. To dissect the molecular function of EWSR1, we successfully established a stable DLD-1 cell line that enables a conditional knockdown of EWSR1 using Auxin Inducible Degron (AID) system. When both *EWSR1* genes of DLD-1 cell were tagged with *mini-AID* at its 5’-end using CRISPR/Cas9 system, treatment of the (*AID-EWSR1/AID-EWSR1*) DLD-1 cells with a plant-based Auxin (AUX) led to the significant levels of degradation of AID-EWSR1 proteins. During anaphase, the *EWSR1* knockdown (AUX+) cells displayed higher incidence of lagging chromosomes compared to the control (AUX-) cells. This defect was proceeded by a lower incidence of the localization of Aurora B at inner centromeres, and by a higher incidence of the protein at kinetochores compared to the control cells during pro/metaphase. Despite these defects, the EWSR1 knockdown cells did not undergo mitotic arrest, suggesting that the cell lacks the error correction mechanism. Significantly, the EWSR1 knockdown (AUX+) cells induced higher incidence of aneuploidy compared to the control (AUX-) cells. Since our previous study demonstrated that EWSR1 interacts with the key mitotic kinase, Aurora B, we generated replacement lines of *EWSR1-mCherry* and *EWSR1:R565A-mCherry* (a mutant that has low affinity for Aurora B) in the (*AID-EWSR1/AID-EWSR1*) DLD-1 cells. The EWSR1-mCherry rescued the high incidence of aneuploidy of EWSR1 knockdown cells, whereas EWSR1-mCherry:R565A failed to rescue the phenotype. Together, we demonstrate that EWSR1 is essential to prevent aneuploidy through interaction with Aurora B, most likely by regulating the localization of Aurora B at centromere.

## INTRODUCTION

The *Ewing sarcoma region 1* gene (*EWSR1*) was originally identified in the pediatric bone cancer, Ewing sarcoma, as a part of an aberrant fusion gene with *FLI1* (Delattre et al., 1992). Subsequent studies showed that *EWSR1* is fused to various types of transcription factors in multiple sarcomas (e.g.*EWSR1-WT1*; desmoplastic small round cell tumor, *EWSR1-ATF1; clear* cell sarcoma, *EWSR1-CHOP*; myxoid liposarcoma, *EWSR1-NR4A3*; extraskeletal myxoid chondrosarcoma) (Panagopoulos et al., 2002, Matsui et al., 2006, Filion et al., 2009, Lee et al., 1997). The EWSR1 has various activities in multiple biological phenomena. For example, EWSR1 interacts with the subunits of TFIID and RNA PolII, and regulates transcription of Oct4 and Brn3a (Zhang et al., 1998, Lee et al., 2005, Gascoyne et al., 2004). The EWSR1 regulates splicing through interaction with splicing factor SF1 as well as small nuclear ribonucleoprotein-specific protein U1C that is required for the early spliceosome formation (Zhang et al., 1998, Knoop and Baker, 2000). Other reports demonstrated that EWSR1 promotes homologous recombination by suppressing R-loop formation (Gorthi et al., 2018). Multiple phenotypes were also reported using animal models with genetically ablated *EWSR1.* The knockout mice for EWSR1 displayed impaired differentiation of pre-B lymphocytes, defects in meiosis of sperm, and reduction of mitochondria through degradation of PGC1a and of mitochondria function in brown fat and skeletal muscles of *EWSR1*-deficient mice (Li et al., 2007, Park et al., 2015). Our laboratory discovered that the zebrafish *ewsr1a* (homologue of human *EWSR1*) mutant displayed impaired differentiation of chondrocytes due to the compromised EWSR1-Sox9 dependent transcription (Merkes et al., 2015). In addition, the zebrafish *ewsr1a* (homologue of human *EWSR1*) mutant displays increased incidence of mitotic dysfunction, aneuploidy, and increased incidence of tumorigenesis in a *tp53* mutation background (Park et al., 2016, Azuma et al., 2007). The study suggested that the loss of EWSR1 may be a part of molecular pathogenesis of EWSR1-expressing sarcomas because its tumor cells lack one EWSR1 allele due to the formation of the fusion gene. One important question that arose from this study is whether loss of EWSR1 leads to aneuploidy after one cell cycle, or if an aneuploid cell population expands by undergoing a selection process. Therefore, this study aimed to elucidate whether loss of EWSR1 in one cell cycle is sufficient to induce aneuploidy, and to characterize the mechanism of the induction of aneuploidy.

A major cause of the induction of aneuploidy is chromosomal mis-segregation derived from defects in microtubule-kinetochore attachment, in altered spindle microtubule dynamics, or in cohesion and condensation of chromosomes during mitosis (Coschi et al., 2010, Lengauer et al., 1997b, Solomon et al., 2011). One form of chromosomal mis-segregation is due to chromosome bridges where a stretched DNA strand connects two segregating chromosomes during anaphase. Chromosome bridges are often derived from dicentric chromosomes, from chromosomes with aberrant condensation/cohesion, or from tangled DNA originating from replication stresses (Hauf et al., 2001, Hetzer, 2010). Other form of chromosomal mis-segregation is derived from lagging chromosomes that are “left behind chromosomes” between segregating chromosomes during anaphase. The lagging chromosomes are often derived from merotelic attachment, an occasion of both microtubules nucleated from opposite centrioles attaching to the same kinetochore of one chromosome (Funk et al., 2016, Cimini et al., 2001b). Lagging chromosomes result in aneuploidy because they are unevenly distributed in two daughter cells, resulting in either gain or loss of the chromosomes (Cimini et al., 2001a). One important regulator of faithful chromosomal segregation is the Aurora B kinase (Aurora B), a component of chromosome passenger complex (CPC) that includes three additional components, INCENP, Borealin and Survivin (Adams et al., 2000, Kaitna et al., 2000). The Aurora B mainly localizes at inner centromeres and kinetochores during pro/metaphase, and later, it is relocated to the spindle midzone (an anti-parallel spindle structure formed between segregating chromosomes) during anaphase (Cimini et al., 2006, Hadders and Lens, 2022, Hümmer and Mayer, 2009, Gruneberg et al., 2004). During pro/metaphase, microtubules are nucleated from the centrioles located at the opposite ends of the cell, and turn over until they attach to the each side of a kinetochore (amphitelic attachment); thus both chromatids are pulled in opposite directions with equal force by microtubules. Aurora B prevents the induction of lagging chromosomes by destabilizing the site of merotelic attachment by phosphorylating Hec1, MCAK and Kif2b. Aurora B also activates the spindle assembly checkpoint (SAC) by enhancing the localization of SAC proteins Mps1 and BubR1 to maintain checkpoint signaling (Gurden et al., 2018) and it also activates the tension checkpoint (also known as error correction) by phosphorylating outer kinetochore proteins such as KMN network (Knl1, Mis12 and Ndc80) (Welburn et al., 2010, DeLuca et al., 2011). In general, when a cell undergoes defects during mitosis, the cell arrests at in prometaphase during the error correction process (Gorbsky, 1997). This coordination prevents the cell from dividing and to mis-segregate its chromosomes without repairing its error.

Our previous study demonstrated that the 565^th^ Arg of EWSR1 that locates in its RGG (Arg-Gly-Gly) 3 domain was the critical amino acid required for the interaction with Aurora B, and it is essential for the localization of Aurora B at the spindle midzone during anaphase (Park et al., 2014). Our following study demonstrated that the *ewsr1a/ ewsr1a* homozygous zebrafish mutant induces mitotic dysfunction, aneuploidy and both *ewsr1a/wt* heterozygous and *ewsr1a/ ewsr1a* homozygous zebrafish promote tumorigenesis in a *tp53/wt* mutant background (Park et al., 2016). In this study, we asked if the function of EWSR1 in preventing aneuploidy is conserved in human cells. We aimed to address whether the loss of EWSR1 induces aneuploidy after one cell cycle, or if aneuploidy is enriched as a result of a selection process after multiple cell divisions. To address these questions, the Auxin Inducible Degron system (AID) was employed in this study as an excellent means to degrade our target protein conditionally. Specifically, the system requires the tagging of the protein with a plant specific ubiquitination sequence, *miniAID* (mAID), within cells expressing OsTIR1 that is a plant specific E3 ligase. Therefore, administration of a plant phytohormone, Auxin (AUX), leads to the conditional degradation of the protein (Natsume et al., 2016). In this study, using CRISPR/Cas9 system, we successfully established a stable line that carries *mAID* tagged *EWSR1* alleles (*AID-EWSR1/AID-EWSR1*) at a homozygous level in OsTIR1 expressing DLD-1 cells (Hassebroek et al., 2020). The DLD-1 (a colorectal cancer) cell line was utilized in this study because it carries a near-diploid karyotype, and has been utilized as a model cell line to study the induction of aneuploidy (Nicholson et al., 2015, Lengauer et al., 1997b). Here, we show that the knockdown of EWSR1 for one cell cycle is sufficient to induce lagging chromosomes and aneuploidy without inducing mitotic arrest. We demonstrate that the impaired EWSR1-Aurora B interaction overrides the error correction mechanism that prevents the induction of aneuploidy.

## METHODS

### Plasmid preparation (DNA construct) and transfection

To establish the (*AID-EWSR1/AID-EWSR1*) DLD-1 cell line, CRISPR/Cas9 system was employed to tag the 5’ end of *EWSR1* gene with *mini-AID (mAID).*

The donor plasmid of *mAID* was generated by following the procedure of the previous report with a minor modification (Hassebroek et al., 2020). First, the homology arms (left/up and right/down) that targets the *EWSR1* locus were amplified from genomic DNA extracted from DLD-1 using following primers respectively.

- *hEWSR1* up HA5’ *PciI:* 5’-CCGACATGTCCGAGAACCCTGGATCCATTCC-3’
- *hEWSR1* up HA3’ *SalI:* 5’-CGTGTGACTTTCTCTCCTTCCTCCTCGTTCTCTC-3’
- *hEWSR1* down HA5’ PAM *SpeI:* 5’ -CCAACTAGTATGGCGTCAACGGGTGAGTATAGTGGAAC-3’
- *hEWSR1* down HA3’ *NotI:* 5’-CAAGCGGCCGCCCCACAAATCCACGGTATCTTTATTTG-3’

Note that mutations were introduced in PAM sequences on the homology arms to avoid the Cas9 dependent degradation of the donor plasmid. The PCR products for the homology arms were inserted into the donor plasmid (containing hygromycin resistant gene/P2A sequence and 3x mini-AID and 3X Flag sequence) using *PciI/SalI* and *SpeI/NotI* sites respectively (Hassebroek et al., 2020).

The guide RNA sequence was designed using the CRISPR Design Tool; https://figshare.com/articles/CRISPR_Design_Tool/1117899 (Rafael Casellas laboratory, National Institutes of Health) and (Zhang laboratory, MIT). The suggested sequences of the guide RNA sequence from these websites are listed below.

- *EWSR1* gRNA1 F: ATGGCGTCCACGGGTGAGTA
- *EWSR1* gRNA1 R: TACTCACCCGTGGACGCCAT
- *EWSR1* gRNA2 F: AGTTCCACCATACTCACCCG
- *EWSR1* gRNA2 R: CGGGTGAGTATGGTGGAACT

The oligos (synthesized by Integrated DNA technologies, IDT) were annealed, inserted into pX330 at its BbsI site (Addgene, #42230), and its sequence was confirmed by the DNA sequence (ACGT Inc.).

The *EWSR1-mCherry* or *EWSR1:R565A-mCherry* DNA constructs that targets the *AAVS1* locus of the (*AID-EWSR1/AID-EWSR1*) DLD-1 cell line are generated as described below. The *mCherry, hEWSR1,hEWSR1:R565A* genes were individually amplified using *pCDNA4-His-maxC-mCherry, pSG5-2XFLAG-hEWSR1* or *pSG5-2XFLAG-hEWSR1:R565A* plasmids as a template using the following primers (Park et al., 2014).

- *MluI-hEWS* F: 5’-GATACGCGTATGGCTGCCACGGATTAC-3’
- *NotInostop3’huEWS1965to1945hEWS* R: 5’-CTGCGGCCGCGTAGGGCCGATCTCTGC-3’
- *NotI mChe* F: 5’ -GCGGCCGCAGGCGCTGG-3’
- *SalI stop mChe* R: 5’-CTTGTCGACTTACTTGTACAGCTCGTCC-3’

The PCR products of *hEWSR1, hEWSR1:R565A* and *mCherry* were inserted to *SalI* and *MluI* sites of pMK243 that was modified at its multicloning sites in the previous study (Tet-OSTIR1-PURO) (Natsume et al., 2016, Hassebroek et al., 2020, Park et al., 2021)

### Establishment and maintenance of the stable cell lines

To establish the (*AID-EWSR1/AID-EWSR1*) DLD-1 cell line, the DLD1-OsTIR cells plated in a 3.5 cm dish was transfected with the *mAID*-donor and guide RNA (gRNA) plasmid constructs using ViaFect (Promega, #E4981) (Mali et al., 2013, Russell C. DeKelver, 2010, Hassebroek et al., 2020). The cells transfected with the DNA constructs were cultured for two days, and split to 10 cm dishes. Next, the cells were further cultured in the medium containing 1μg/mL Blasticidin (Invivogen, #antbl) and 200μg/mL Hygromycin B Gold (Invivogen, #ant-hg) until they form colonies for ten to fourteen days. Sixteen colonies were isolated, and the integration of transgenes were verified with PCR using the genomic DNA obtained from each clone using the following primers;

- *hEWSR1* GenPCR 5’ F: 5’ -CCCGGGTACTCACTGCACGAG-3’
- *hEWSR1* GenPCR 3’ R: 5’-CGGCTTGGGGCTGGAAGC-3’

Among sixteen clones, we chose clone #19, a homozygous (*AID-EWSR1/AID-EWSR1*) clone, for the following analysis (referred to as OsTIR7-19 in the following section). Conditional degradation of mAID tagged EWSR1 protein in the AUX treated OsTIR7-19 cells was confirmed by Immunocytochemistry and Western blot analysis using anti-FLAG (Sigma, #F7425) and anti-EWSR1 (Santa Cruz, #sc-48404) respectively.

To establish the (*AID-EWSR1/AID-EWSR1;EWSR1-mCherry*) and (*AID-EWSR1/AID-EWSR1;EWSR1:R565A-mCherry*) DLD-1 cell lines, Tet-On transgene that contains *EWSR1-mCherry* or *EWSR1:R565A-mCherry* was integrated into the safe harbor *AAVS1* locus of the OsTIR7-19 (*AID-EWSR1/AID-EWSR1*) DLD-1 cell line. The OsTIR7-19 cells were plated in a 3.5 dish, and transfected with the donor and guide RNA plasmids targeting *AAVS1* locus (Addgene, AAVS1 T2 CRISPR in pX330, #72833) (described in the previous section) using ViaFect™ (Promega, #E4981). The cells were cultured for 2 days, re-plated in 10 cm dishes, and were selected for ten to fourteen days with a selection medium (contains 1 μg/ml Blasticidin [Invivogen, #ant-bl], 200 μg/ml Hygromycin B gold [Invivogen, #ant-hg] and 1μg/mL Puromycin). Twenty four clones were screened for the mCherry signal visually with fluorescent microscope, and the integration of transgene in its genome was confirmed by genomic PCR using following primers.

- *AAVS1* F: 5’ -CTGCCGTCTCTCTCCTGAGT-3’
- Pause Site R: 5’ -GTTTTGATGGAGAGCGTATGTTAGTAC-3’
- *SV40* F: 5’ -CCGAGATCTCTCTAGAGGATCTTTGTGAAG-3’
- *AAVS1* R: 5’-CAAAAGGCAGCCTGGTAGAC-3’

The concentration of antibiotics used in this study are 1 ug/ml Puromycin (PUR^R^), Invivogen), 1 μg/ml Blasticidin (BSD^R^) (Invivogen, #ant-bl) and 200 μg/ml Hygromycin B (HYG^R^) gold (Invivogen, #ant-hg). The (*AID-EWSR1/AID-EWSR1*) cells were maintained in the McCoy’s 5A medium that contains BSD and HYG. The (*AID-EWSR1/AID-EWSR1;EWSR1-mCherry*) and (*AID-EWSR1/AID-EWSR1;EWSR1:R565A-mCherry*) DLD-1 cells were maintained in the McCoy’s 5A medium with BSD, HYG and PUR.

To knockdown the AID-EWSR1, the cells were treated with 500 μM indole-3-acetic acid (3-IAA, or AUX) for 24 hr. To express exogenous EWSR1-mCherry or EWSR1:R565A-mCherry, both cell lines were treated with 1 μg/mL of doxycycline (DOX) for 24 hr.

The expression of EWSR1-mCherry and EWSR1:R565A-mCherry proteins were confirmed by immunocytochemistry using rat anti-RFP (1:1000 dilution) (Bulldog Bio Inc, #RMA5F8) followed by anti-Rat Alexa Fluor 568 (1:500 dilution) (Invitrogen, #A-11077), and by western blotting using chicken anti-mCherry antibody (1:1,000 dilution, LSBio, #C-204825), followed by IRDye 800 CW donkey anti-chicken IgG secondary antibody (1:10,000 dilution) (LI-COR, #926-32218).

### Western blotting

Whole cell lysates obtained from cells with/without AUX/DOX were subjected to western blotting. Primary antibodies used in this study are; mouse anti-C9 EWS antibody (1:1000 dilution) (Santa Cruz, #sc-48404); rabbit anti-FLAG antibody (1:1000 dilution) (Sigma, #F7425); rabbit anti-CyclinB antibody (1:1000 dilution, Sigma, #C8831); mouse anti-β-actin antibody (1:1000 dilution, Sigma, #A2228); chicken anti-mCherry antibody (1:1000 dilution, LSBio, #C-204825). Secondary antibodies used in this study are; IRDye 680RD donkey anti-mouse IgG (1:10,000 dilution, LI-COR, #926–68072), IRDye 800CW donkey anti-rabbit IgG (1:10,000 dilution, LI-COR, #926-32213), IRDye 800CW donkey anti-mouse IgG (1:10,000 dilution, LI-COR, # 926-32212), IRDye 800CW donkey anti-chicken IgG (1:10,000 dilution, LI-COR, # 926-32218). All images of western blotting were captured by LI-COR Odyssey Imaging System.

### Chromosome spread

The cells grown in a T25 flasks at ~70-80% confluency were synchronized to mitosis using the thymidine/nocodazole protocol (Kotýnková et al., 2016, Park et al., 2021). The mitotic cells were collected in a 15 ml tube, washed for 3 times with McCoy’s 5A media (without FBS), and it were resuspended in 1 mL of PBS. Half (500 μL) of the mitotic cell suspension was transferred to new tube by mixing with 1 mL of dH_2_O, and it was incubated for 5 min at room temperature. Then, the mitotic cell suspension solution (250 μL) was loaded into a cytology funnel (BMP, #CYTODB25), and it was subjected to cytospin for 5 min at 1000 rpm with max acceleration (Thermo Shandon Cytospin3 Centrifuge, #TH-CYTO3). The slides were subjected to immunocytochemistry as described in the following section.

### Immunocytochemistry

The mitotic cells were subjected to immunocytochemistry by following the protocol described in the previous report (Park et al., 2014). The antibodies used in this study are; Mouse anti-C9EWS antibody (1:500 dilution) (Santa Cruz, #sc-48404); rabbit anti-FLAG antibody (1:500 dilution)(Sigma; rabbit anti-Aurora B antibody(1:500 dilution)(abcam,#ab2254); guinea pig anti-CENPC antibody (1:500 dilution)(MBL, #PD030); mouse anti-α-tubulin antibody (1:4,000 dilution)(Sigma, #T8328); anti-mouse Alexa fluor 568 (1:500 dilution)(Invitrogen,# A-11004); Rat anti-RFP (1:1,000 dilution) (Bulldog Bio Inc, #RMA5F8), anti-rabbit Alexa fluor 488 (1:500 dilution)(Invitrogen, #A32790);anti-guineapig Alexa fluor 568 (1:500 dilution)(Invitrogen, #A-21450) anti-rat Alexa Fluor 568 (1:500 dilution) (Invitrogen, #A-11077).

The chromosomes spread on a slide was fixed with 100 uL of 4% Paraformaldehyde (PFA) at room temperature for 5 min, washed for three times with 0.01% Triton X-100/Phosphate Buffered Saline (PBS) for 5min at room temperature respectively, permeabilized with ice-cold methanol for 5 min at -20°C, and washed for three times with PBS for 5 min at room temperature. Then, the chromosomes on the slide was treated with 100ul of blocking solution (1% Fetal Bovine Serum (FBS)/PBS) at room temperature for 30 min. The slides were subjected to immunocytochemistry with following primary antibodies; rabbit-anti-Aurora B antibody (1:500 dilution) (Abcam, #ab2254) guineapig-anti CENPC antibody (1:500 dilution) (MBL, #PD030), rat-anti RFP (1:500 dilution) (Bulldog Bio Inc, #RMA5F8) for over-night at 4°C. Next day, the chromosomes spread on the slide were washed with PBS at room temperature for 5 min by repeating three times, and incubated with following secondary antibodies; anti-Rabbit Alexa Fluor 488 (1:500 dilution) (Invitrogen, # A32790), anti-guineapig Alexa Fluor 647 (1:500 dilution) (Invitrogen # A-21450), or anti-rat Alexa Fluor 568 (1:500 dilution) (Invitrogen, #A-11077) for 1hr at room temperature, washed with PBS at room temperature for 5 min by repeating three times, and mounted on the VECTASHILED Antifade Mounting Medium with DAPI (Vector laboratory, #H-1200).

### Aneuploidy analysis

The cells were plated at ~30% confluency, followed by the treatment with/without AUX/DOX for 48 hours. Concurrently, the cells were treated with thymidine/nocodazole, and the mitotic cells were subjected to cytospin as described in the previous section. Mitotic chromosomes were visualized with following antibodies; rabbit-Topo2A antibody (gift from Dr. Yoshiaki Azuma) followed by Alexa flour 488 anti-rabbit secondary antibody; rat-RFP antibody followed by Alexa flour 568 anti-rat secondary antibody (Invitrogen); anti-guineapig CENP-C antibody (MBL) followed by Alexa Flour 647 anti-guineapig antibody (Invitrogen). All antibodies were diluted at 1:500 dilution rate.

### Image documentation

All images of cells and chromosomes were acquired using the Nikon Plan Apo 60x or 100×/1.4 oil objective lens on a TE2000-U microscope (Nikon) with a Retiga SRV CCD camera (QImaging) operated by Meta Morph imaging software (MetaMorph Inc.) at room temperature.

### Statistical analysis

All graphs are presented as mean with standard deviation (S.D) or standard error of the mean (S.E.M). The statistical analysis was conducted with one-way or two-way ANOVA followed by Tukey’s multiple comparison test or two-tailed paired t-test using GraphPad9 software (confidence was defined at *p* < 0.05).

## RESULTS

### The EWSR1 regulates faithful chromosomal segregation

To determine whether loss of EWSR1 induces aneuploidy after one cell cycle, or cells with aneuploid expand during a long period of time, it is essential to control the timing of the knockdown of EWSR1. To accomplish this, we employed the Auxin-Inducible Degron (AID) system to degrade EWSR1 conditionally in DLD-1 cells (Yesbolatova et al., 2019). The major reason for utilizing this conditional knockdown system is that it allows us to assess the effect of EWSR1 within a single cell cycle, thus the phenotypic change caused by the knockdown of protein of interest leads to a better understanding of the activity of the protein. The CRISPR/Cas9 DNA constructs, a *mAID-3xFLAG-Hyg^r^* (Hyg, Hygromycin resistant gene) and two guide RNA constructs that target the start codon of the *EWSR1* gene, were transfected into the stable DLD-1 cell line that expresses a plant derived E3 ligase*, OsTIR1* (integrated at a *RCC1* locus) (**Fig 1A**) (Hassebroek et al., 2020). The DLD-1 cell line was used in this study because it has a near-diploid karyotype, and it has been used to model the activity of proteins of interest in aneuploidy induction (Lengauer et al., 1997a). The cells transfected with the CRISPR/Cas9 DNA constructs were selected with hygromycin, and the genomic DNAs of the colonies were amplified by PCR to verify the integration of the *mAID-3xFLAG-Hyg^r^* construct. Among a total of 48 colonies that were screened, 11 clones carried the *AID-EWSR1 wt* genotype, and 5 clones carried the *AID-EWSR1 AID-EWSR1* genotype (**Fig S1**). Among the *AID-EWSR1 AID-EWSR1* homozygous clones, we chose clone #19 to assess the function of EWSR1 in the following study.

**Fig 1.**
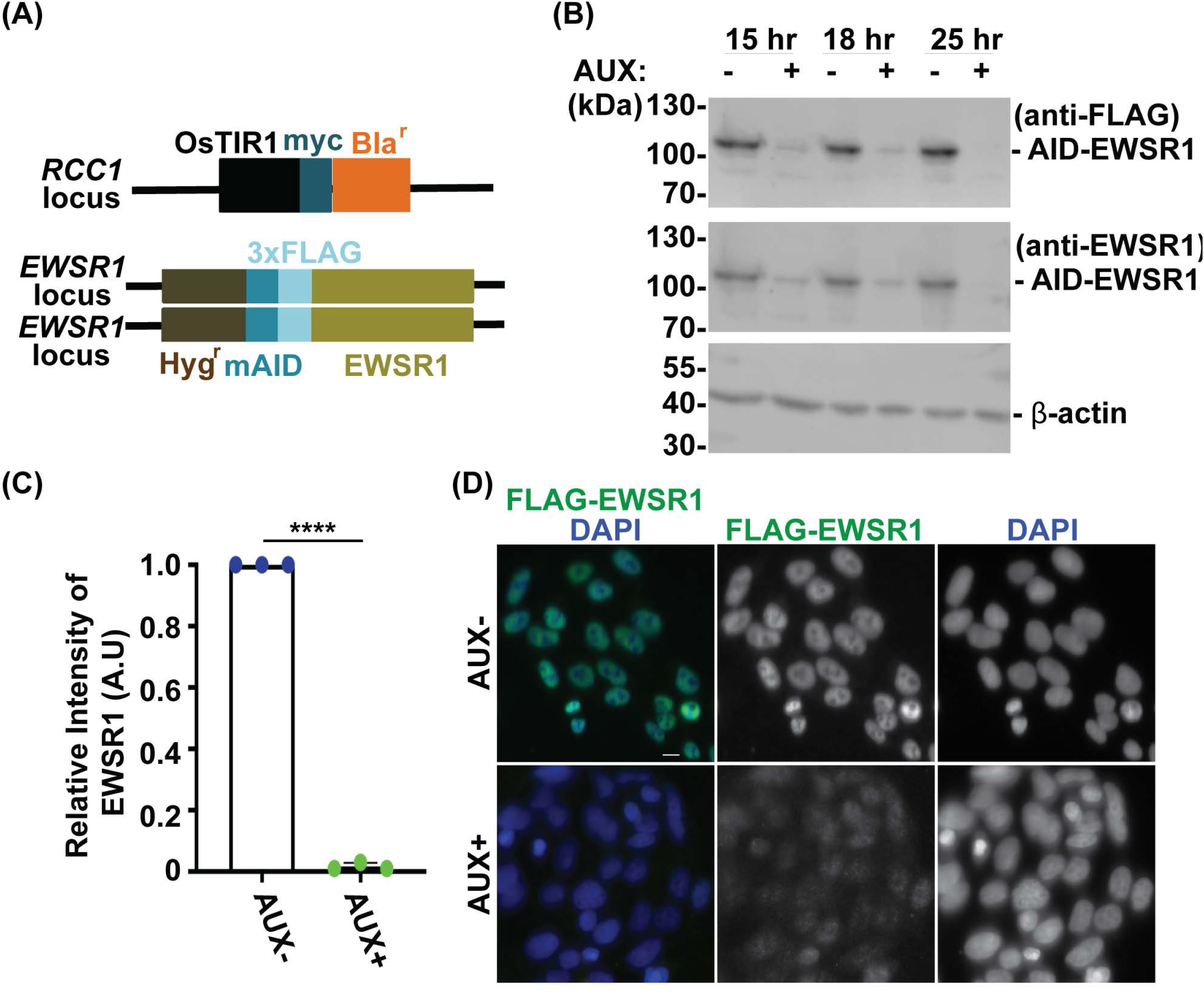
The treatment of *AID-EWSR1/AID-EWSR1* stable DLD-1 cell line with Auxin induces an efficient degradation of AID-EWS. **A.** Schematic diagram of *OsTIR1* and *AID-EWSR1* DNA construct. *OsTIR1*: plant driven E3 ligase, *Bla^r^:* Blasticidine resistance gene, *Hyg^r^:* Hygromycin resistance gene, *mAID:* mini Auxin-Inducible Degron tag, *3XFLAG;* three tandem repeat of FLAG tag. The *OsTIR1* construct was inserted to *RCC1* locus, and *mAID* construct was inserted to both *EWSR1* loci (homozygous). **B.** Representative images of western blot using anti-FLAG (top panel), anti-EWSR1 (middle panel) and anti-β actin (bottom panel) obtained from AUX- and AUX+ cells in the presence of Auxin with given time point (15 hours, 18 hours and 25 hours). **C.** Relative intensity of the bands of western blotting visualized with anti-EWSR1 (normalized to the bands with anti-β-actin) obtained from AUX- and AUX+ cells (n=3 experiments). Graph shows the mean of each group with Standard Deviation (SD). (n = 3 experiments). \*\*\*\**p* < 0.0001 (Two tailed paired *t*-test). **D.** Representative images of immunocytochemistry using anti-FLAG (green) and DAPI (blue) obtained from AUX- and AUX+ cells. Scale bar= 10um.

We began our study by characterizing the Auxin dependent degradation of EWSR1 in the newly established (*AID-EWSR1/AID-EWSR1*) DLD-1 cell line. Because the division time of DLD-1 cells is approximately 22 hrs, the cells were treated with 500 μM of Auxin (AUX+) for 15, 18 and 25 hrs, and it was subjected to western blotting using anti-FLAG and anti-EWSR1 antibodies to assess the level of its degradation (Dexter et al., 1979). The cells treated with AUX for 25 hrs accomplished maximum degradation of EWSR1 (**Fig 1B**). Repetition of the experiment three times revealed a consistent reduction of the signal of EWSR1 protein (normalized by β-actin) in the AUX+ sample when it was compared to the AUX-sample (****p<0.0001) (**Fig 1C**). Based on this result, the cells were treated with AUX for a minimum of 24 hrs in the following experiments. In addition, immunocytochemistry using anti-FLAG antibody confirmed drastic reduction of the FLAG-EWSR1 protein in the AUX treated (AUX+) cells (**Fig 1D**). Together, treatment of *AID-EWSR1 AID-EWSR1* cells with AUX leads to efficient degradation of EWSR1.

We previously identified *EWSR1a and EWSR1b* in zebrafish (the homologues of human *EWSR1*), and proposed that the proteins regulate faithful chromosomal segregation during anaphase (Azuma et al., 2007). The analysis was done in 24 hrs post-fertilization (hpf) zebrafish, thus the study did not address whether the defects were directly derived from a single cell cycle, or from secondary defects derived from multiple cell cycles. To elucidate this question, the stable cell line was treated with AUX, and the incidence of aberrant chromosome segregation in the EWSR1 knockdown (AUX+) cells was compared to the control (AUX-) cells. Both AUX- and AUX+ cells were first synchronized in mitosis using the conventional thymidine/nocodazole protocol, and the samples were fixed at 60 min after the release from nocodazole **(Fig 2A**). The chromosomes of AUX- and AUX+ cells were visualized with DAPI, and the incidence of aberrant segregation of chromosomes, lagging chromosomes and chromosome bridges, was scored. The AUX+ cells displayed significantly higher incidence of lagging chromosome compared to the AUX-cells (**Fig 2B and C**). Contrary, there was no significant change of the incidence of chromosome bridges between AUX- and AUX+ cells (n=3 experiments) (**Fig 2B and C**). Note that the incidence of chromosome bridges in AUX+ cells was slightly increased compared to AUX-cells. Together, the results suggests that EWSR1 maintains chromosomal stability by preventing the induction of lagging chromosomes.

**Fig 2.**
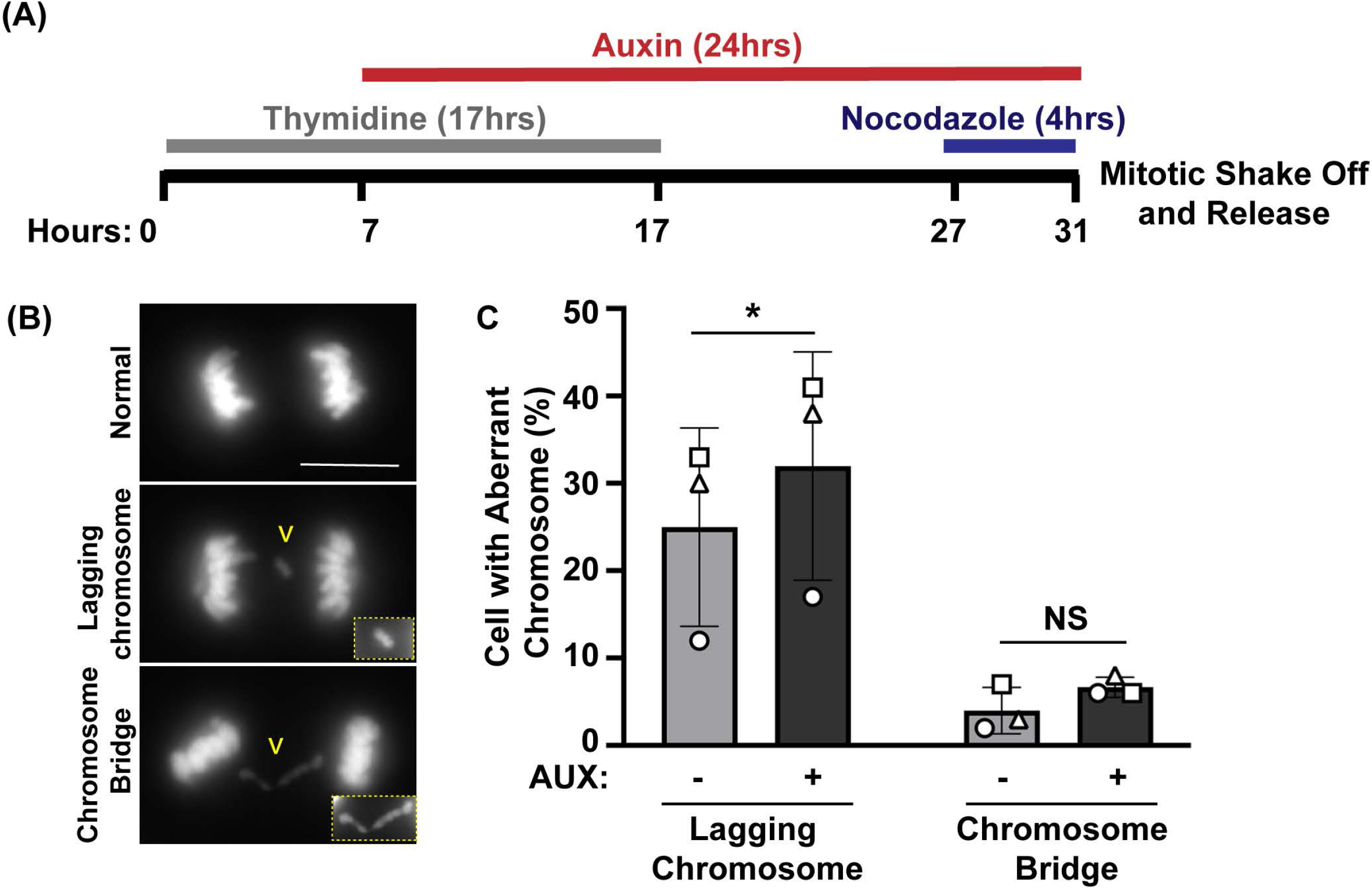
The EWSR1 knockdown promotes induction of lagging chromosome during anaphase. **A.** Schematic diagram for the mitotic synchronization of EWSR1 knockdown cells using Thymidine and Nocodazole, followed by the mitotic shake-off and release from the mitotic arrest for 60 min. **B.** Representative images of chromosomes; normal (top panel), lagging chromosome (middle panel) and chromosome bridge (bottom panel). Yellow square highlights the magnified images of either lagging chromosome or chromosome bridge. Scale bar= 10um. **C.** Percentages of lagging chromosomes (left graph) and of chromosomal bridges (right graph) in AUX- and AUX+ cells (Total 34-104 anaphase cells per sample, n=3 experiments). \**p*<0.05. (paired t-test), NS; Non-Significant.

### The EWSR1 protein is required for the proper localization of Aurora B at inner centromeres

To accomplish faithful chromosome segregation, microtubules nucleated from one side of centrosome has to attach to the same side of the kinetochore. If microtubules attach to both/same kinetochores but only from one centrosome (syntelic attachment) or if most attachments are correctly oriented but some microtubules from the opposite side of centrosome connect to the same kinetochores (merotelic attachment), the result will be lagging chromosomes during anaphase (Cimini et al., 2001a). During the error correction process, Aurora B kinase, the enzymatic subunit of the chromosome passenger complex (CPC) is recruited to the site of error (Ohi et al., 2003). Therefore, we examined whether the localization of Aurora B is altered in the EWSR1 knockdown (AUX+) cells. The cells were synchronized at prometaphase using thymidine/nocodazole treatment, released for 30 min, followed by the cytospin procedure to spread the cells on a slide. Then, mitotic chromosomes on the slides were subjected to immunocytochemistry using anti-Aurora B (green) and a centromeric marker, anti-CENP C (red). The localization of Aurora B was scored based on four criteria; inner centromere, kinetochore proximal centromere (KPC), both inner centromere and KPC and no signal (**Fig. 3A**).

**Fig 3.**
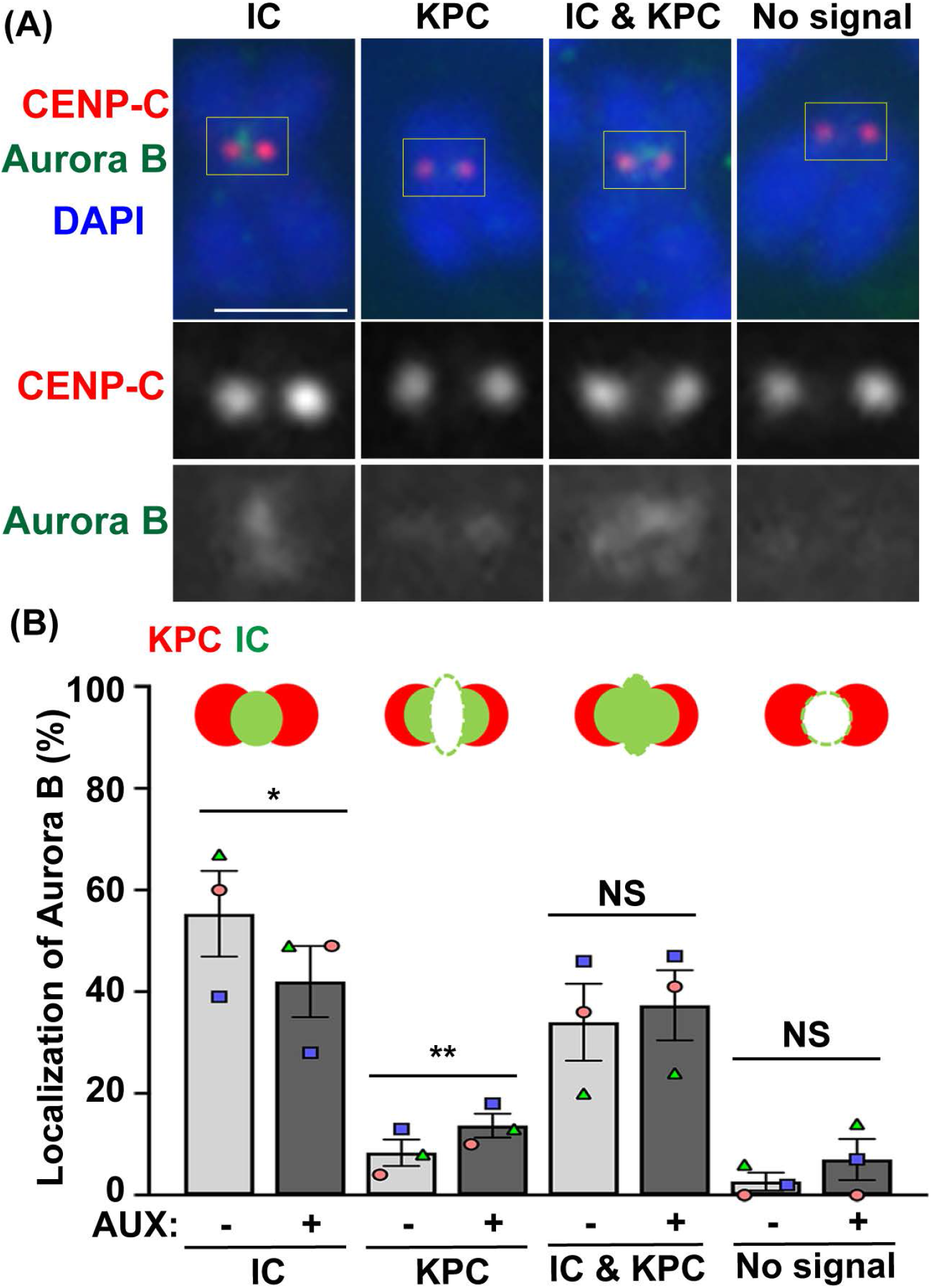
The EWSR1 is required for Aurora B localization at inner centromere, and prevents its localization at kinetochore during early mitosis. **A.** Representative images of Aurora B on pro/metaphase chromosome obtained from control (AUX-) cells that were released from thymidine/nocodazole for 30 min. The Aurora B (Green, top and bottom panel) and CENPC (kinetochore marker) (Red, top and middle panel) were visualized by immunocytochemistry using anti-Aurora B and anti-CENPC antibodies. Classification of the localization of Aurora B; Inner Centromere (IC), Kinetochore Proximal Centromere (KPC), Inner Centromere and Kinetochore Proximal Centromere (IC+KPC), and No Signal. Scale bar= 10um. **B.** Quantification of Aurora B localization at IC, KPC, IC+KPC, and No signal between AUX- and AUX+ cells (total of 712-732 chromosomes per sample, n=3 experiments). Values are mean with standard deviation (S.D.). Two-tailed paired t-test; \**p*<0.05, \*\**p*<0.01, NS; Non-Significant.

There was a significantly decreased incidence of Aurora B at inner centromeres, and an increased incidence of the localization of Aurora B at KPC in the EWSR1 knockdown cells (AUX+) compared to those of control cells (AUX-) (**Fig. 3B**). The data suggests that EWSR1 is required for the proper localization of Aurora B at the inner centromere. This may occur in response to improper microtubule-kinetochore interactions and could be related to the increased anaphase lagging chromosomes.

### The EWSR1 knockdown cell does not undergo mitotic arrest

In general, when the process of chromosomal segregation is impaired, the cell should be arrested at mitosis so that the error is corrected. Cells are arrested before anaphase onset when microtubules are erroneously connected to kinetochores, or before cytokinesis when chromosomes lag or bridge during anaphase. Because the EWSR1 knockdown (AUX+) cells displayed high incidence of lagging chromosomes, we aimed to determine whether the EWSR1 knockdown (AUX+) cells underwent mitotic arrest. The cells were treated with/without Auxin (AUX- and AUX+), and the cells were synchronized in mitosis using the same thymidine/nocodazole protocol described in **Fig 2A**. The cells arrested at prometaphase were harvested (0 min), released for 30 min, 60 min and 90 min, and it were plated on a coverslip. The stages of mitosis were scored based on the shape of chromosomes (DAPI signal), and its incidences were compared between AUX- and AUX+ cells. To our surprise, there were no significant differences between the duration of the mitosis of AUX- and AUX+ cells (**Fig 4A**).

**Fig 4.**
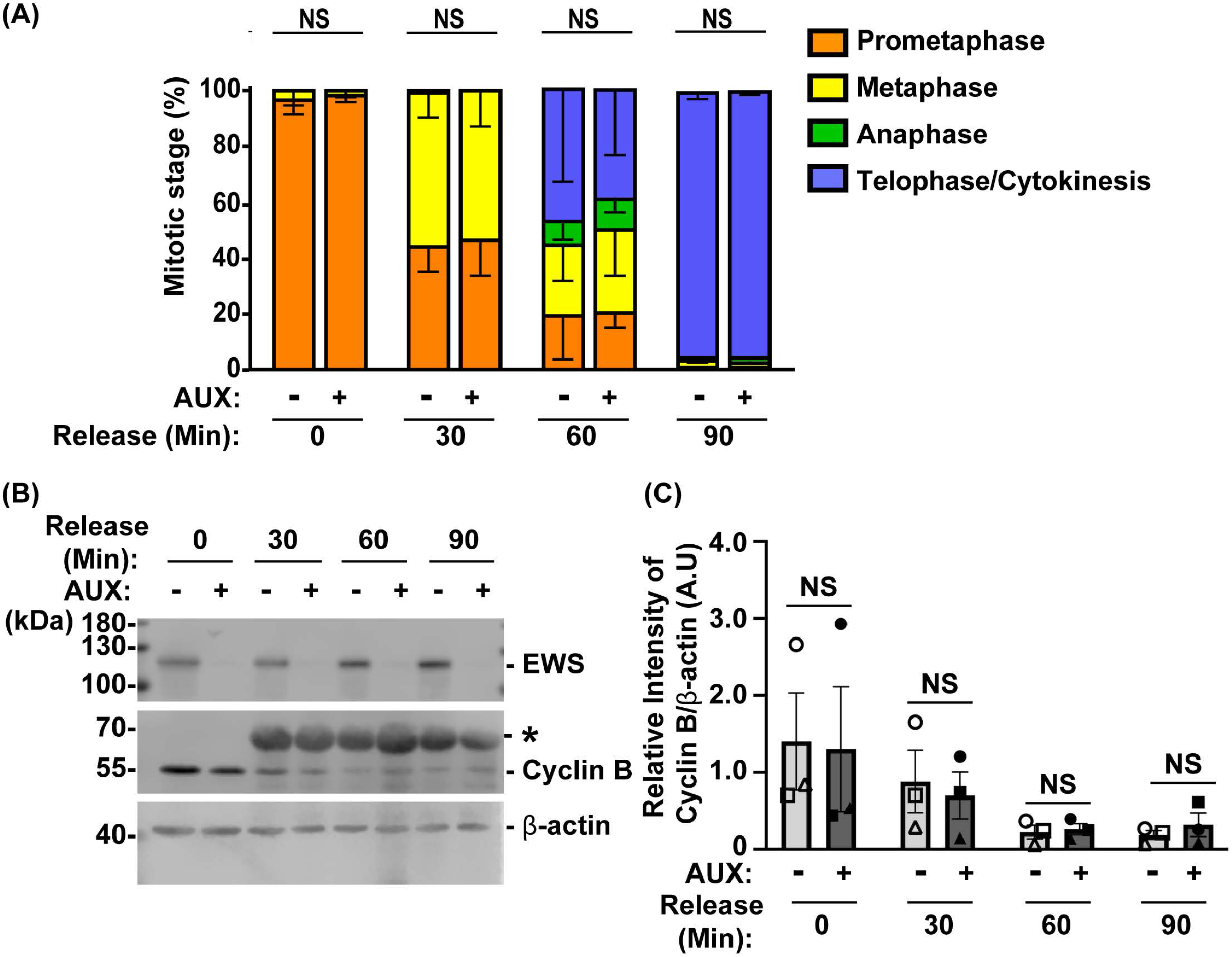
The EWSR1 knockdown cells do not arrest at any mitotic stages. **A.** The percentages of mitotic stage of AUX- and AUX+ cells released for 0, 30, 60 and 90 min after the thymidine/nocodazole treatment (total of 83-173 cells per sample, n=3 experiments). NS=non-significant. **B.** Representative images of western blot using anti-FLAG (top panel), anti-Cyclin B (middle panel) and anti-ß actin (bottom panel) obtained from AUX- and AUX+ cells released at 0, 30, 60, and 90 min after the thymidine/nocodazole treatment. *: Non-specific band. **C.** Quantification of the levels of Cyclin B protein (normalized by β-actin) obtained from AUX- and AUX+ cells (n=3 experiments). Images of western blotting gel using anti-EWSR1 (top panel), anti-Cyclin B (middle panel), and anti-β-actin antibodies (bottom panel). Values are mean with S.D. Two-way ANOVA with Tukey’s multiple comparison test. NS: Non-Significant.

To further verify the result, we employed a biochemical approach to quantify the level of Cyclin B, a protein that is known to undergo degradation during anaphase (Minshull et al., 1990). The results showed that there was no significant difference in the level of Cyclin B proteins between AUX- and AUX+ cells in all time course samples (**Fig 4B and 4C**). Despite that the EWSR1 is required for faithful chromosomal segregation, the results suggest that EWSR1 knockdown does not affect the duration of mitosis by activating checkpoint controls.

### The EWSR1 protein prevents the induction of aneuploidy through interaction with Aurora B

Because the EWSR1 knockdown (AUX+) cells displayed higher incidence of lagging chromosomes compared to the control (AUX-) cells within a single cell cycle (**Fig 2**), we further examined whether the AUX+ cells result in the induction of aneuploidy after one cell division. The cells were treated with Auxin for 48 hrs combined with cell synchronization because this protocol allowed us to efficiently deplete EWSR1, and then quantify ploidy after one mitosis (**Fig 5A**). Samples were spread onto slides using a cytospin to prepare chromosome spreads, and were subjected to immunocytochemistry using anti-CENPC (red, a marker for centromere) or anti-TOPO2A (green, a marker to visualize the chromosomes). The representative images of the chromosomes from two sample groups are shown in **Fig 5B**. When the numbers of chromosomes in both sample groups were scored (43-50 cells per sample, n=3 experiments), the EWSR1 knockdown (AUX+) cells displayed higher incidence of aberrant numbers of chromosomes compared to the control (AUX-) cells (**Fig 5C**, and **Fig 5D**; the percentages of the chromosomal number in both sample groups). Together, the results suggest that the EWSR1 prevents the induction of aneuploidy within a single cell division.

**Fig 5.**
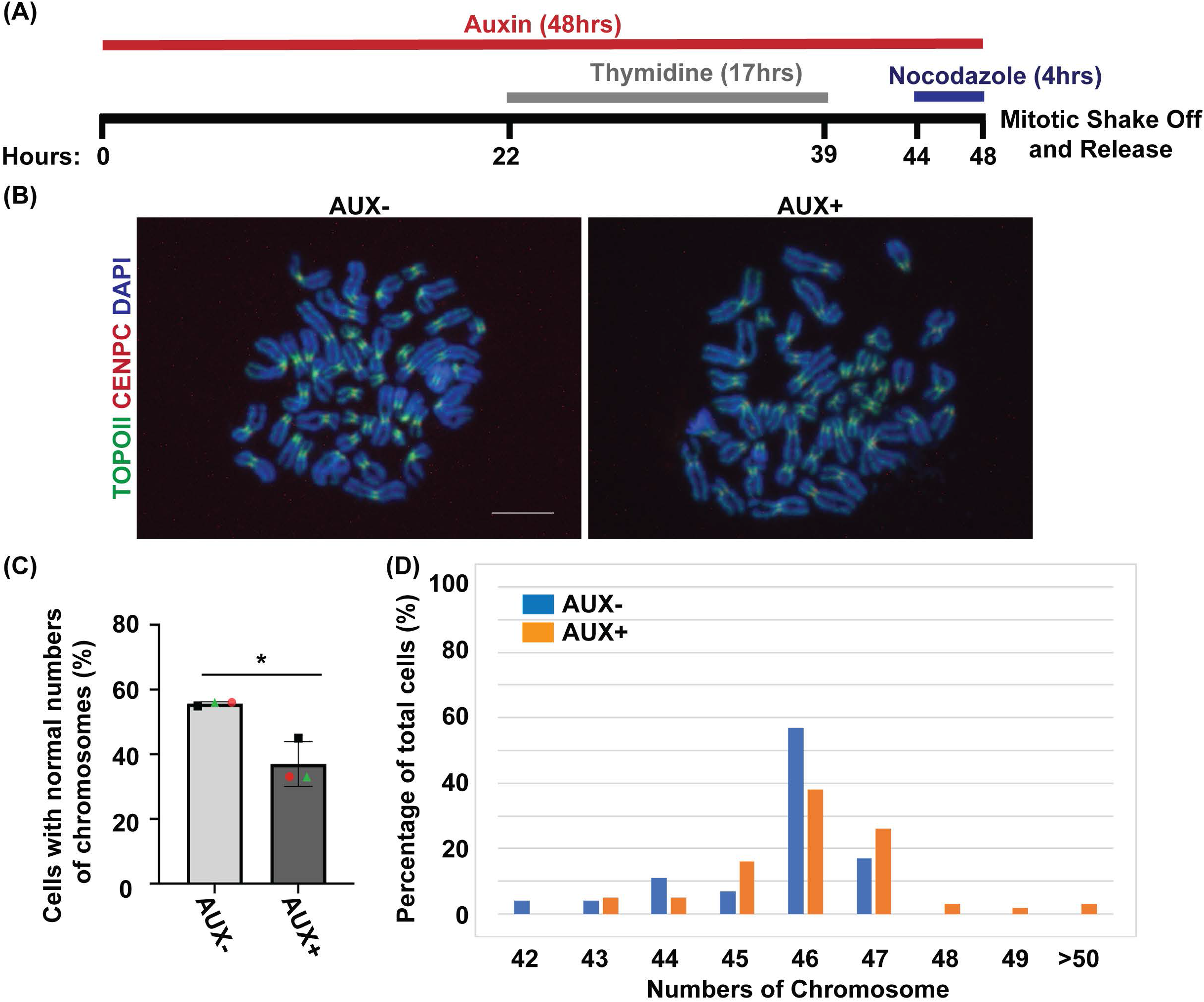
The EWSR1 knockdown promotes the induction of aneuploidy. **A.** Schematic diagram for the mitotic synchronization of AID-EWSR1 knockdown (synchronized to mitosis using thymidine/nocodazole, and treated with AUX for 48 hrs concurrently) cells. The mitotic cells were harvested 30 min after released from the arrest. **B.** Representative images of chromosomes visualized with anti-CENPC (red), and anti-Topoisomerase II (green) obtained from AUX- and AUX+ cells. Scale bar= 10um. **C.** The percentage of cells that displayed normal 46 numbers of chromosomes was decreased in AUX+ cells compared to AUX-cells. 18-24 cells per sample were counted from n=3 experiments (Total of n=56 cells from AUX-, and of n=62 cells from AUX+). Two-tailed paired t-test, *p<0.05. **D.** The percentage of cells for each of the chromosome numbers in AUX- and AUX+ cells. The percentages were calculated from total chromosomes obtained from n=3 experiments.

Our previous study showed EWSR1 interacts with Aurora B, whereas EWSR1:R565A, a mutant of 565th Arg of EWSR1 substituted to Ala, displayed a reduced level of the interaction (Park et al., 2016). To evaluate whether EWSR1-Aurora B interaction is required for preventing the induction of aneuploidy, we employed the CRISPR/Cas9 system to generate two new replacement stable cell lines that have an integration of Tet-on *EWSR1-mCherry (mCherry* tag is fused to the C-terminus of *EWSR1),* and *EWSR1:R565A-mCherry* constructs at a safe harbor *AAVS1* locus of the DLD-1 (*AID-EWSR1/AID-EWSR1*) cell line. The CRISPR/Cas9 DNA constructs were transfected into DLD-1 (*AID-EWSR1/AID-EWSR1*) cells, and the colonies were selected with puromycin (**Fig S2**). Then, the presence of mCherry signals in the colonies were visually screened. Among the twenty four colonies, two mCherry positive colonies for both *EWSR1-mCherry* and *EWSR1:R565A-mCherry* transfected cells were isolated respectively. The cells were treated with 500 μM of Auxin (AUX+), and 1μg/mL of doxycycline (DOX) for 24hrs. The knockdown of EWSR1 and expression of the exogenous EWSR1-mCherry or EWSR1:R565A-mCherry mutant within each of the cell lines were verified with immunocytochemistry, and with western blot using anti-FLAG and anti-mCherry antibodies obtained from cells (**Fig S3 and S4**).

To evaluate whether the interaction between EWSR1 and Aurora B is required for the induction of aneuploidy, both (*AID-EWSR1/AID-EWSR1; EWSR1-mCherry*) and (*AID-EWSR1/AID-EWSR1; EWSR1:R565A-mCherry*) DLD-1 cells were treated with combinations of AUX and DOX (AUX-/DOX-, AUX+/DOX-, and AUX+/DOX+) for 48 hrs, and the chromosomes were spread onto slides using a cytospin, followed by immunocytochemistry using anti-TOPO2A and anti-CENPC. The treatment of the (*AID-EWSR1/AID-EWSR1; EWSR1-mCherry*) cells with AUX induced a high incidence of aneuploidy compared to non-treated (AUX-/DOX-) cells, whereas EWSR1 knockdown/ EWSR1-mCherry expressing (AUX+/DOX+) cells rescued the high incidence of aneuploidy (**Fig 6A**). Percentages of chromosome numbers per cell are listed in **Fig 6B**. The treatment of the (*AID-EWSR1/AID-EWSR1; EWSR1:R565A-mCherry*) cells with AUX consistently induced high incidence of aneuploidy compared to non-treated (AUX-/DOX-) cells; however, the EWSR1 knockdown/EWSR1:R565A-mCherry expressing (AUX+/DOX+) cells did not rescue the high incidence of aneuploidy (**Fig 6C and D**). These results suggest that the interaction between EWSR1 and Aurora B is required for the prevention of aneuploidy.

**Fig 6.**
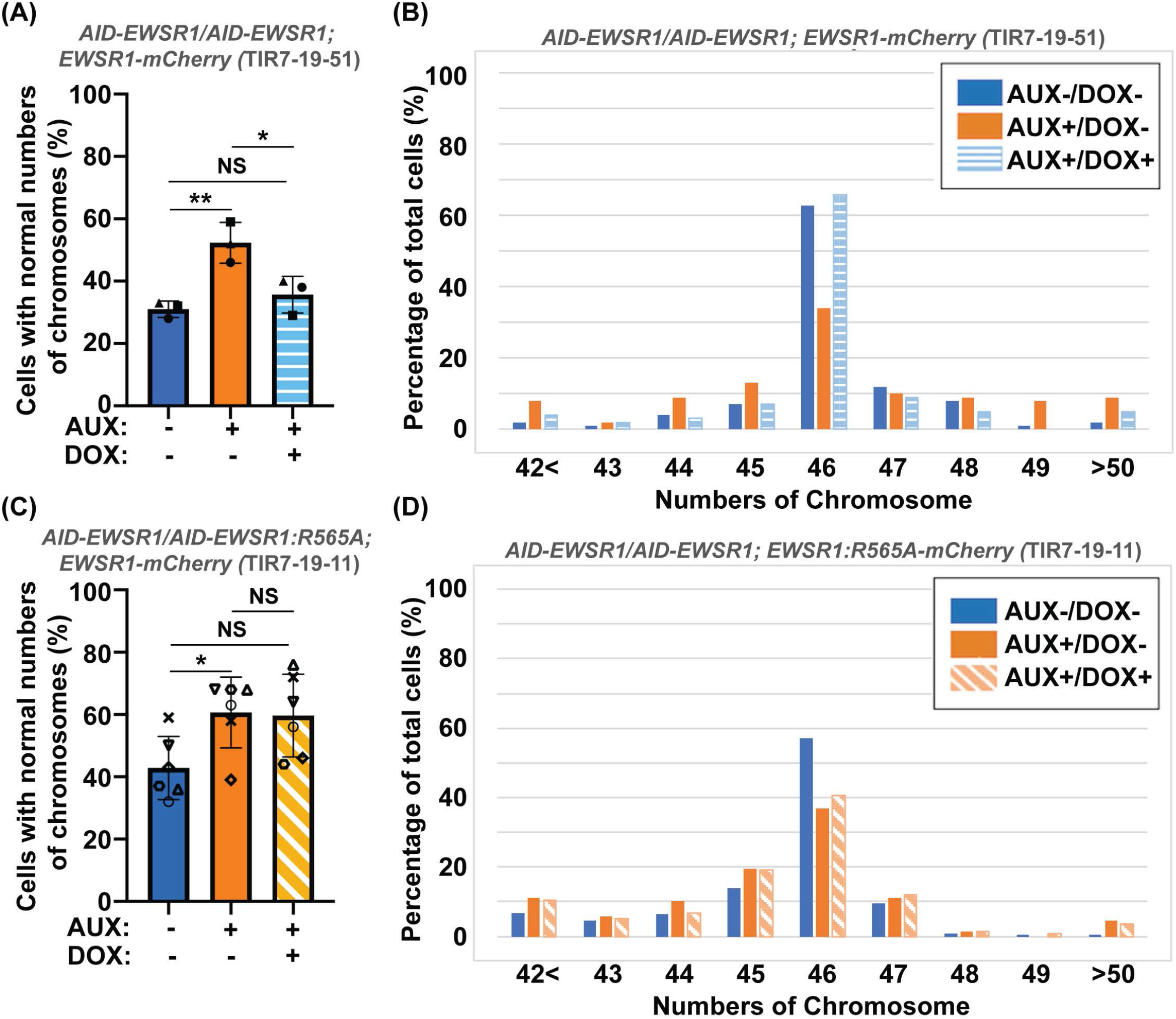
The interaction between EWSR1 and Aurora B prevents the induction of aneuploidy. **A.** The high incidence of aberrant chromosome numbers in AUX+/DOX-was rescued by the expression of EWSR1 overexpression (AUX+/DOX+) cells (28-59 cells per sample, n=3 experiments). One-way ANOVA with Tukey multiple comparison test., **p<0.01, *p<0.05. NS=Non-significant. **B.** The percentages of the chromosome numbers in AUX-/DOX-(n=122), AUX+/DOX- (n=117), and AUX+/DOX+(n=106) cells. **C.** High incidence of aberrant chromosome numbers in AUX+/DOX-cells was not rescued by the expression of EWSR1:R565A in AUX+/DOX+ cells. (35-42 cells per sample, n = 6 experiments, One-way ANOVA with Tukey multiple comparison test. *p<0.05, NS=non-significant. **D.** The percentage of the numbers of chromosome in AUX-/DOX- (n=221), AUX+/DOX- (n=228), and AUX+/DOX+(n=219) cells.

## DISCUSSION

The aim of this study was to establish if depletion of EWSR1 protein in human cells causes aneuploidy and, if so, to elucidate the molecular function of EWSR1 in preventing the induction of aneuploidy. Here, we demonstrate that the conditional knockdown of EWSR1 in DLD-1 cells for one cell cycle is sufficient to evict Aurora B from inner centromere and to enrich the protein at KPC during pro/metaphase, to induce high incidence of lagging chromosomes during anaphase, and to induce high incidence of aneuploidy after one mitosis. Note that this is the first study to show the EWSR1 knockdown dependent aneuploidy induction in human cells. Importantly, we propose that EWSR1 inhibits the induction of aneuploidy by interacting with Aurora B. Despite that the EWSR1 knockdown cells induced lagging chromosomes and aneuploidy during mitosis, the cells failed to undergo mitotic arrest. Our study highlights a potential new role of EWSR1-Aurora B in error correction during mitosis, and cells lacking EWSR1 may induce chromosomal instability (CIN) by overriding the process of error correction (**Fig 7**).

**Fig 7.**
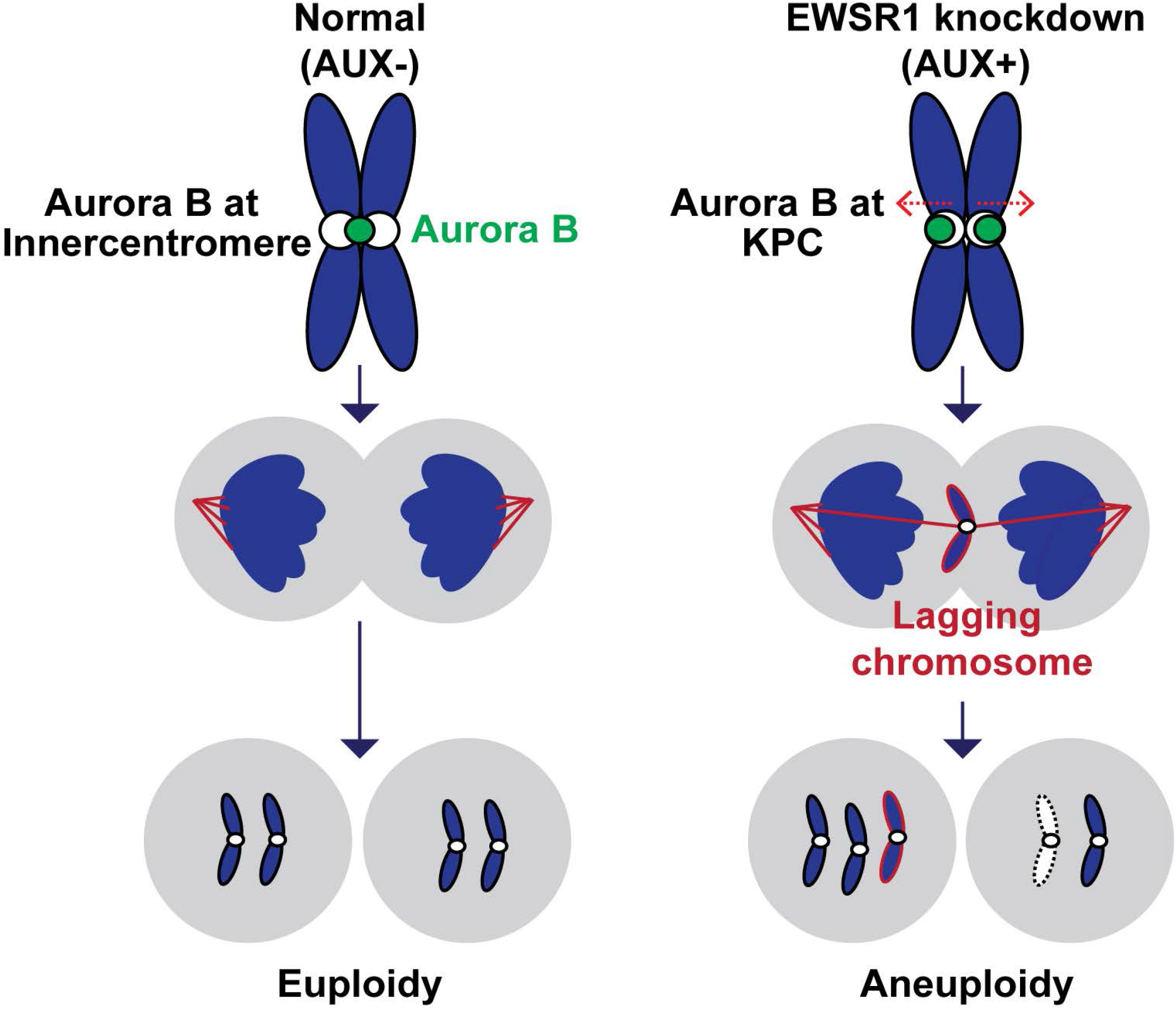
Schematic model of the study. The EWSR1 knockdown leads to the relocation of Aurora B from inner centromere to KPC (prometaphase and metaphase), induction of lagging chromosomes (anaphase), and induction of aneuploidy (after a single mitosis). The EWSR1 prevents the induction of aneuploidy through interaction with Aurora B. Despite of the induction of chromosome mis-segregation in EWSR1 knockdown cells, the cells do not arrest at mitosis, and it is likely that the cells are overriding the error collection process.

Future studies are required to elucidate how EWSR1 prevents the induction of lagging chromosomes and aneuploidy, and how EWSR1 knockdown cells escape from mitotic arrest. In general, microtubule turnover occurs until it establishes a proper bi-oriented microtubule-kinetochore attachment during prometaphase. When there is an attachment error, Aurora B is directly recruited to the kinetochore, where it corrects the error by detaching the microtubule from the kinetochore (Broad et al., 2020, Caldas et al., 2013). Therefore, one possibility for the function of EWSR1 is that the protein promotes detachment of microtubules by regulating the kinase activity of Aurora B at the erroneously attached kinetochore. For example, EWSR1 may be required for the Aurora B dependent phosphorylation of its substrates, MCAK and Kif2b (microtubule depolymerizing proteins) because both proteins inhibit the induction of chromosome mis-segregation by promoting the microtubule turnover (Wordeman et al., 2007). Other possibility is that EWSR1 facilitates the Aurora B dependent phosphorylation of Hec1/Ndc80, a core element of kinetochores that activates the detachment of microtubules from kinetochores and promotes polymerization of the microtubule at its plus end (DeLuca et al., 2006). Another potential mechanism required for faithful chromosome segregation is that EWSR1 regulates R-loops at centromeres and resolves lagging chromosome by activating the ATR-Aurora B axis (Kabeche et al., 2018). One attractive model is that EWSR1 functions as an adapter molecule to recruit molecules listed above, and to coordinate the two pathways to resolve the error to prevent the induction of lagging chromosomes. Lagging chromosomes are often induced by merotelic attachment, a condition where a kinetochore is attached to microtubules nucleated from two opposite centrosomes (Cimini et al., 2001a). It is noteworthy that our assay system to measure the lagging chromosomes employed the mitotic synchronization protocol using thymidine/nocodazole. The nocodazole induces higher incidence of merotelic attachment, thus the nocodazole treatment in the EWSR1 cells may have enabled us to observe the pronounced activity of EWSR1 preventing the induction of lagging chromosome (Cimini et al., 2001a).

There are high numbers of fusion genes that contain the N-terminus of *EWSR1* expressed in various sarcomas, leukemias and a carcinoma (Endo et al., 2016, Panza et al., 2021, Panagopoulos et al., 2002, Matsui et al., 2006, Filion et al., 2009). All *EWSR1* fusion expressing cells retains one wildtype *EWSR1* allele due to the formation of a single fusion gene. In support of this, our previous study using zebrafish demonstrated that the loss of one *ewsa* (a homologue of human *EWSR1*) allele (*ewsa/wt*) promotes tumorigenesis in the *tp53* mutation background, it is highly possible that haploinsufficiency of EWSR1 contributes to tumorigenesis in the *EWSR1* fusion expressing tumor cells. Our study provides a strong platform to address the effect of loss of one *EWSR1* allele in the induction of lagging chromosomes, aneuploidy, and the activity of Aurora B by generating the *EWSR1* fusion tumor models that carries (*EWSR1/wt;EWSR1* fusion) genotypes in future studies. For this reason, knowledge of the molecular function of EWSR1 may shed light to the *EWSR1* fusion expressing sarcomas.

## ACKNOWLEDGMENTS

This study was supported by the grants R03CA223949, P30CA168524, University of Kansas General Research Fund (GRF), Midwest Cancer Alliance (MCA) Advisory Board Funding, and Bionexus KC Patton Trust Research Grants. We would like to thank Dr. Duncan Clark (University of Minnesota) for his critical comments for the manuscript.

## DATA AVAIRABILITY

The data presented in this study has been deposited to BioRXiV. https://www.biorxiv.org/content/10.1101/2022.06.17.496636v1

