## Supplementary material for "EWSR1 prevents the induction of aneuploidy by regulating the localization of Aurora B at inner centromere": SUPPLIMENTAL INFORMATION

### Supplementary Materials

**Figure S1. Establishment of *AID-EWSR1/AID-EWSR1* in DLD1 cell by CRISPR/Cas9 genome editing system.**

**A.** Schematic diagram of the donor plasmid. LHA: left homology arm of *EWSR1*, RHA: right homology arm of *EWSR1*, p2A: self-cleaving peptide, 3xmAID: three-repeat of mini Auxin-Inducible Degron, 3XFLAG; three-repeat of FLAG tag. Red arrows: PCR primers to verify the integration of the donor construct in the clone. **B**. Agarose gel images of the PCR amplicons that verify the integration of the donor construct at the *EWSR1* locus. (Homozygous: 2.8kbp, heterozygous: 2.8kbp and 1.2kbp, No integration:1.2 kbp bands). The clone #19 (homozygous) was utilized in this study. PC: positive control (amplified from the genome amplified from the parental wildtype cell), NC: negative control (dw).

**Fig S2. Establishment of *AID-EWSR1/AID-EWSR1;EWSR1-mCherry* or *AID-EWSR1/AID-EWSR1; EWSR1:R565A-mCherry* using CRISPR/Cas9 genome editing system.**

Schematic diagram of the donor plasmid for integrating the *EWSR1-mCherry* and *AAVS1-EWSR:R565A-mCherry* to the *AAVS1* locus of (*AID-EWSR1/AID-EWSR1*) DLD-1 cell; short homology arm sequences of *AAVS1* (LHA/RHA)-Tet-On 3G-gene of interest (*EWS-mCh/EWSR565A-mCh)* -PuroR targeting safe harbor locus *AAVS1*. LHA: left homology arm; RHA: right homology arm; Tet-On 3G: expressing rtTA protein that binds Tet-On promoter in the presence of doxycycline; PuroR: Puromycin resistant gene tag. Blue/Purple arrows indicate primer sets for confirmation of the integration of gene cassette.

**Fig S3. The treatment of the (*AID-EWSR1 AID-EWSR1;EWSR1-mCherry)* DLD-1 cell line enables efficient degradation of AID-EWSR1 and ectopic expression of EWSR1-mCherry.**

**A.** Representative images of immunocytochemistry for expression of EWSR1-mCherry visualized with anti-mCherry (red) and of AID-EWSR1 visualized with anti-FLAG (green) obtained from the cells treated with/without AUX/DOX (AUX-/DOX-, AUX-/DOX+, AUX+/DOX- and AUX+/DOX+) for 24hrs. Merged images (left panel), anti-mCherry (second from left panel), anti-FLAG (second from right panel), and DAPI (right panel). Scale bar= 10um. **B.** Representative images of western blotting using anti-FLAG (Top panel), anti-mCherry (middle panel), and anti-β actin (bottom panel) obtained from the cells with AUX-/DOX-, AUX-/DOX+, AUX+/DOX- and AUX+/DOX+ (treated for 24hrs). *: non-specific band. **C.** Relative intensity of the bands of anti-mCherry (normalized to bands of anti-β-actin) obtained from western blotting. **D.** Relative intensity of the bands of anti-FLAG (normalized to bands of anti-β-actin) obtained from western blotting. Graph shows the mean of each group with Standard Deviation (SD) (obtained from n = 3 experiments).

**Fig S4. The treatment of the (*AID-EWSR1/ AID-EWSR1;EWSR1:R565A-mCherry)* DLD-1 cell line enables efficient degradation of AID-EWSR1 and expression of EWSR1;R565A-mCherry.**

**A.** Representative images of the cells treated with/without AUX/DOX (treated for 24hrs), followed by the visualization of the expression of EWSR1:R565A-mCherry with anti-mCherry (red) and of AID-EWSR1 visualized with anti-FLAG (green) by immunocytochemistry. Merged images (left panel), anti-mCherry (second from left panel), anti-FLAG (second from right panel), and DAPI (right panel). Scale bar= 10um. **B.** Images of western blotting visualizing AID-EWSR1using anti-FLAG (Top panel), EWSR1:R565A-mCherry using anti-mCherry (middle panel), and of β-actin using anti-β actin (bottom panel) from the cells with AUX-/DOX-, AUX-/DOX+, AUX+/DOX- and AUX+/DOX+ treated for 24hrs. *: non-specific band. **C.** Normalized intensity of western blotting bands of anti-mCherry that were normalized with the bands of anti-β-actin. **D.** Normalized intensity of western blotting bands of anti-FLAG, normalized to bands obtained from the usage of anti-β-actin. Graph shows the mean of each group with Standard Deviation (SD) (obtained from n = 3 experiments).

**Fig S5. The expression of EWSR1-mCheery rescues the induction of aneuploidy in the EWSR1 knockdown cells, whereas EWSR1:R565A-mCheery lacks the activity.**

Representative images of chromosomes visualized with anti-CENPC (red), anti-Topoisomerase II (green), and DAPI (blue) obtained from (*AID-EWSR1/AID-EWSR1;EWSR1-mCherry*) cells (**A**), and (*AID-EWSR1/AID-EWSR1;EWSR1:R565A-mCherry*) cells (**B**). Scale bar= 20um.

Fig S1

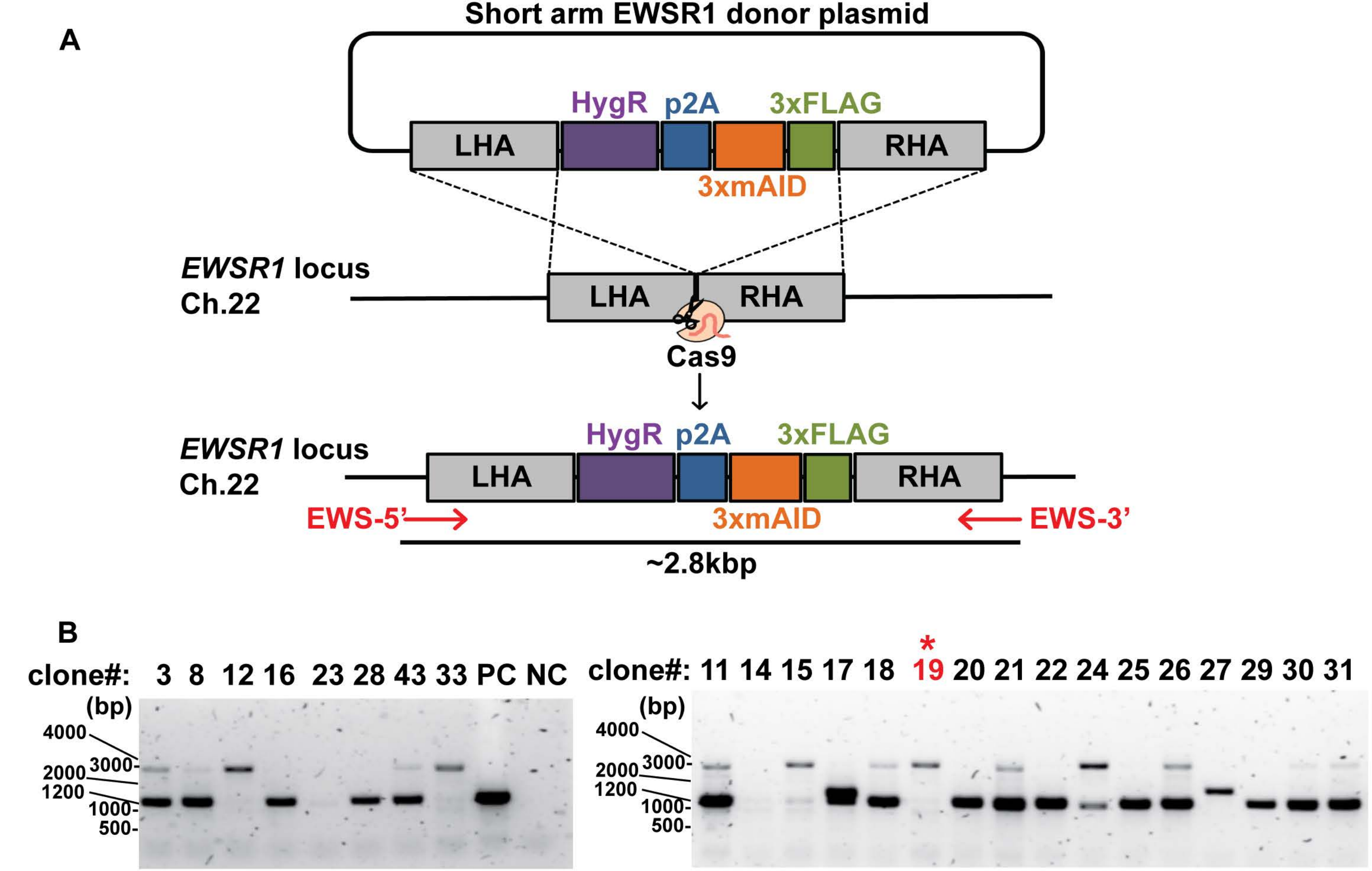
A
Short arm EWSR1 donor plasmid
HygR
p2A
3xFLAG
LHA
RHA
3xmAID
EWSR1 locus
Ch.22
LHA
RHA
Cas9
EWSR1 locus
Ch.22
HygR
p2A
3xFLAG
LHA
RHA
EWS-5'
3xmAID
EWS-3'
~2.8kbp
B
clone#: 3 8 12 16 23 28 43 33 PC NC
(bp)
4000
3000-
2000
1200
1000-
500-
*
clone#: 11 14 15 17 18 19 20 21 22 24 25 26 27 29 30 31
(bp)
4000
3000-
2000
1200
1000-
500-

FigS2

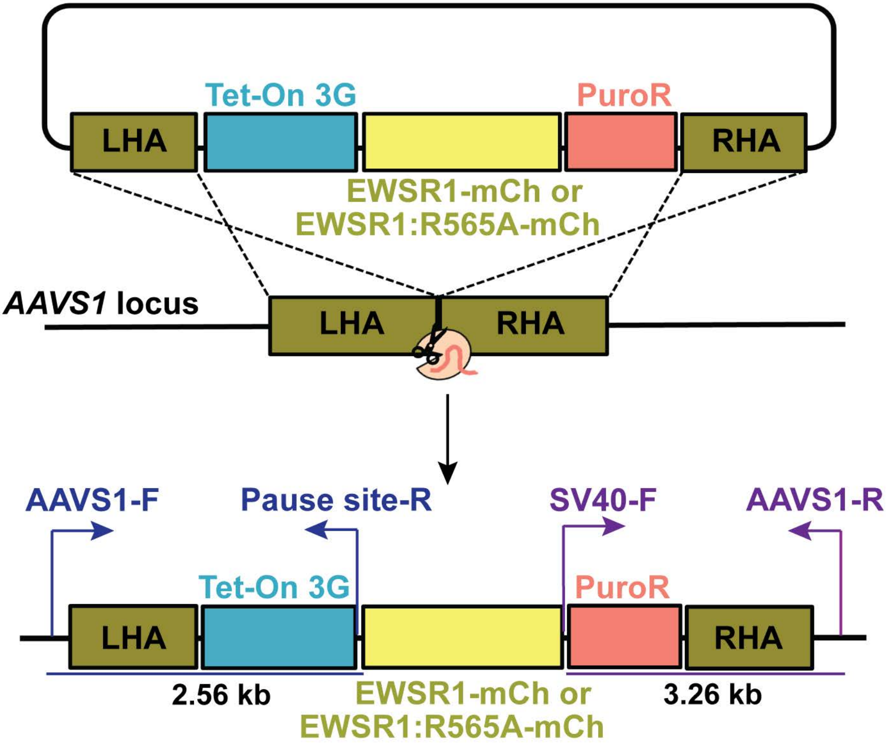
Tet-On 3G
PuroR
LHA
RHA
EWSR1-mCh or
EWSR1:R565A-mCh
AAVS1 locus
LHA
RHA
AAVS1-F
Pause site-R
SV40-F
AAVS1-R
Tet-On 3G
PuroR
LHA
RHA
2.56 kb
EWSR1-mCh or
EWSR1:R565A-mCh
3.26 kb

Fig S3

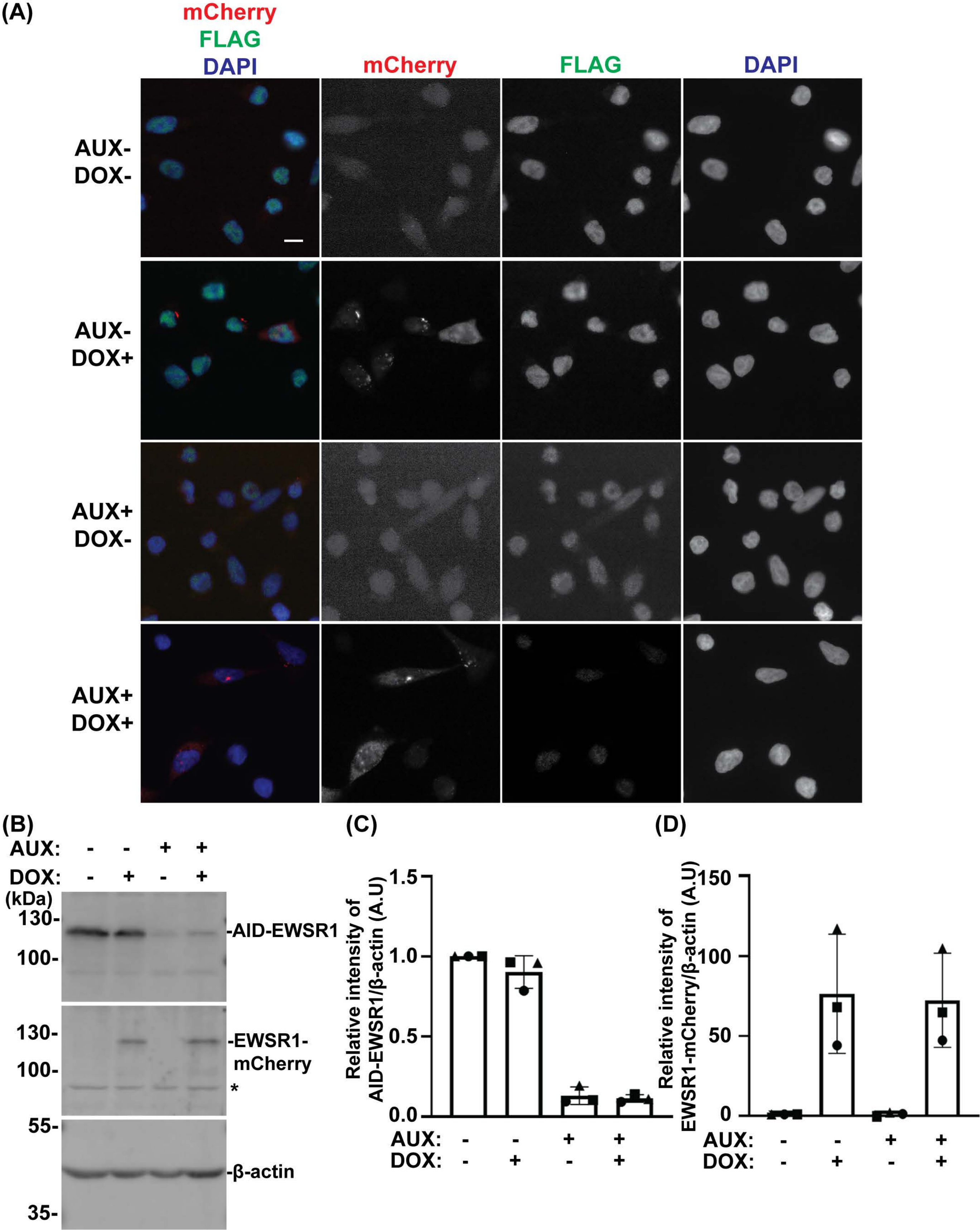
(A)
mCherry
FLAG
DAPI
mCherry
FLAG
DAPI
AUX-
DOX-
AUX-
DOX+
AUX+
DOX-
AUX+
DOX+
(B)
AUX: - - + +
DOX: - + - +
(kDa)
130-
100-
-AID-EWSR1
130-
100-
-EWSR1-
mCherry
*
55-
-β-actin
35-
(C)
Relative intensity of
AID-EWSR1/β-actin (A.U)
1.5
1.0
0.5
0.0
AUX: - - + +
DOX: - + - +
(D)
Relative intensity of
EWSR1-mCherry/β-actin (A.U)
150
100
50
0
AUX: - - + +
DOX: - + - +

Fig S4

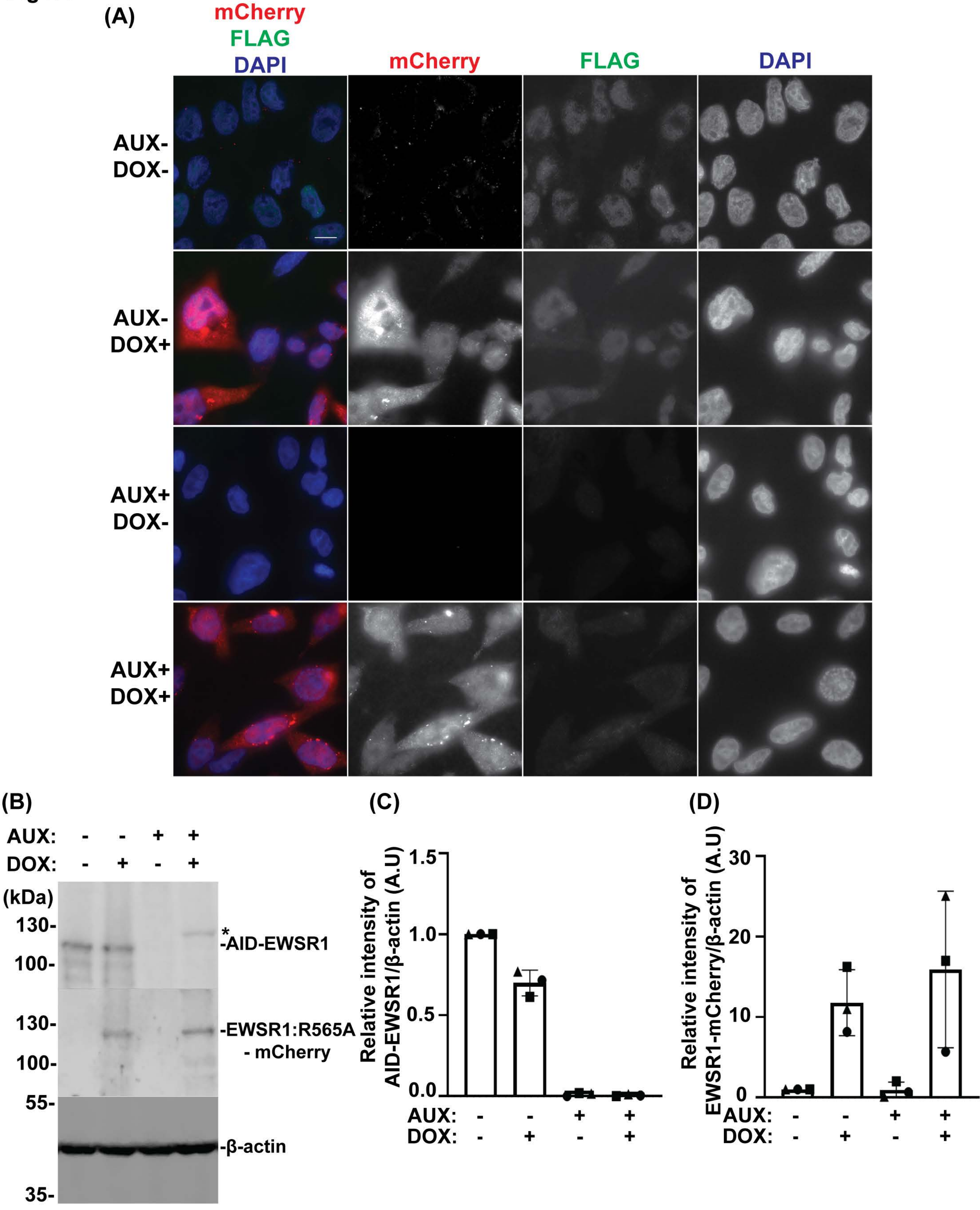

(A)
mCherry
FLAG
DAPI
mCherry
FLAG
DAPI
AUX-
DOX-
AUX-
DOX+
AUX+
DOX-
AUX+
DOX+
(B)
AUX: - - + +
DOX: - + - +
(kDa)
130-
100-
130-
100-
55-
35-
*
-AID-EWSR1
-EWSR1:R565A
- mCherry
-β-actin
(C)
Relative intensity of
AID-EWSR1/β-actin (A.U)
1.5
1.0
0.5
0.0
AUX: - - + +
DOX: - + - +
(D)
Relative intensity of
EWSR1-mCherry/β-actin (A.U)
30
20
10
0
AUX: - - + +
DOX: - + - +

FIGS5

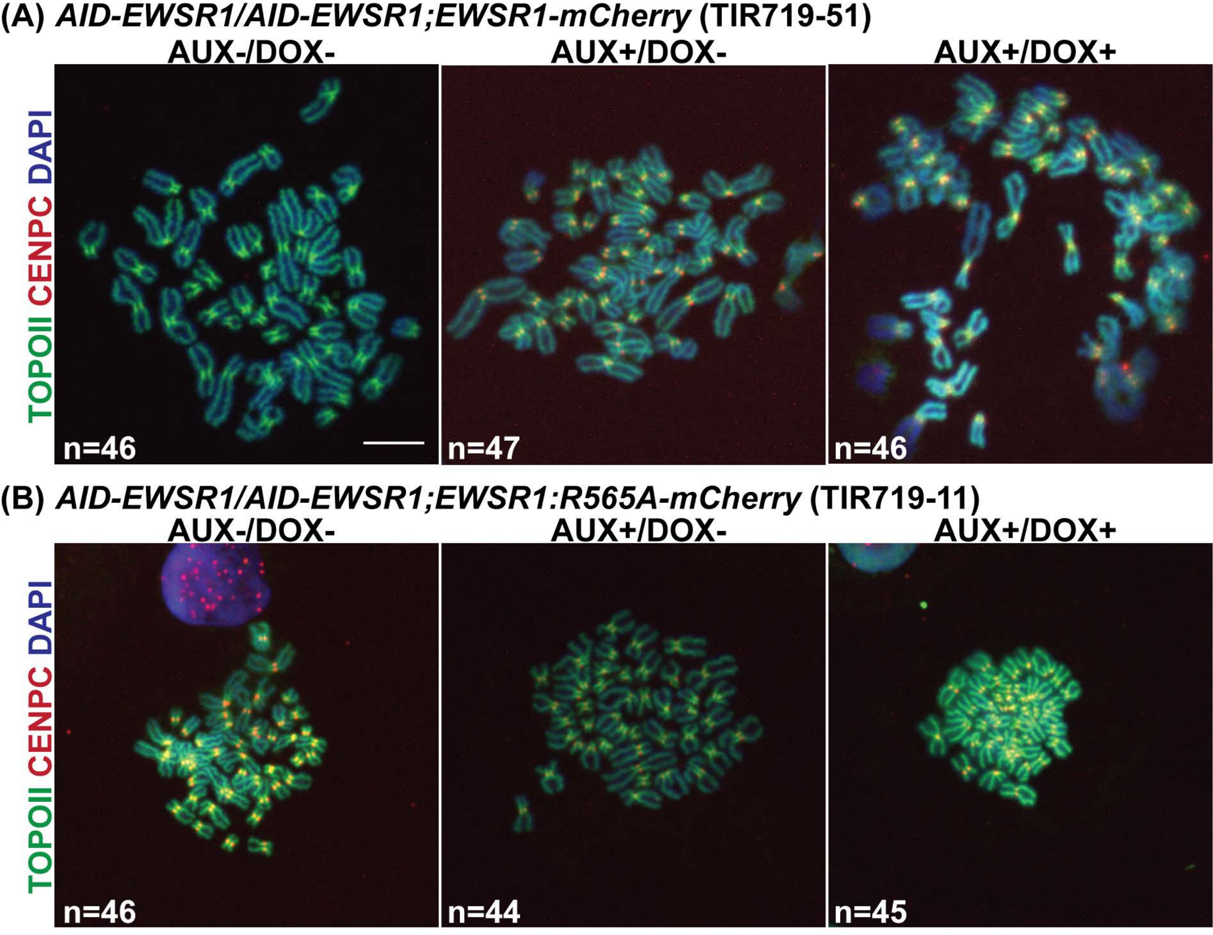
(A) AID-EWSR1/AID-EWSR1;EWSR1-mCherry (TIR719-51)
AUX-/DOX-
AUX+/DOX-
AUX+/DOX+
TOPOII CENPC DAPI
n=46
n=47
n=46
(B) AID-EWSR1/AID-EWSR1;EWSR1:R565A-mCherry (TIR719-11)
AUX-/DOX-
AUX+/DOX-
AUX+/DOX+
TOPOII CENPC DAPI
n=46
n=44
n=45
